# Discovery of cell-permeable inhibitors for human succinyl-CoA:glutarate-CoA transferase

**DOI:** 10.64898/2026.11.12.765678

**Authors:** Frank H. Rizzo, Tetyana Dodatko, Brandon Stauffer, Chunli Yu, Kunal Kumar, Robert J. DeVita, Michael B. Lazarus, Sander M. Houten

## Abstract

Glutaric Aciduria Type I (GA-1) is a recessive, serious neurometabolic disorder characterized by a build-up of toxic metabolites in the lysine metabolic pathway with no current pharmacological treatment. Previous mouse studies have shown that a deficiency in SUGCT attenuates the accumulation of some of these metabolites *in vivo*. These results highlight the potential for SUGCT as a therapeutic target in treating GA-1. We had previously identified biochemical inhibitors for SUGCT and had solved the crystal structure of the protein, both apo and bound to an inhibitor. However, we were unable to demonstrate cellular activity for these compounds. In this study, we report findings from additional *in vitro* high-throughput screening and address these challenges by developing an improved cell-based assay to detect SUGCT inhibition. We show that by improving the permeability of valsartan using its ethyl ester prodrug, SUGCT is effectively inhibited in GA-1 cells. We also identify meclofenamic acid as a new SUGCT inhibitor with cellular activity, providing additional validation and highlighting this enzyme target potential for GA-1.

## INTRODUCTION

Succinyl-CoA:glutarate-CoA transferase (SUGCT) is a mitochondrial enzyme mainly expressed in the liver and kidney that belongs to the type-III CoA transferase (CaiB/BaiF) superfamily of proteins.^1–3^ Its main physiological function is catalyzing the succinyl-CoA-dependent re-esterification of free glutarate into glutaryl-CoA, which is considered an example of metabolite repair (Figure 1, S1A).^1,3–5^ Without this repair, glutarate, which is derived from lysine, hydroxylysine, and tryptophan catabolism, would be a dead-end metabolite that is excreted in urine.^7^ Glutaric Aciduria Type III (GA-3) is a benign, biochemical phenotype caused by loss-of-function variants in *SUGCT* (formerly *C7orf10*).^2,6^ While GA-3 is a biochemical abnormality with low or no clinical significance, our previous work using mouse models has demonstrated that a naturally occurring *Sugct* variant is a biochemical modifier of the Glutaric Aciduria Type I (GA-1) phenotype (Figure S2).^7^

**Figure 1:**
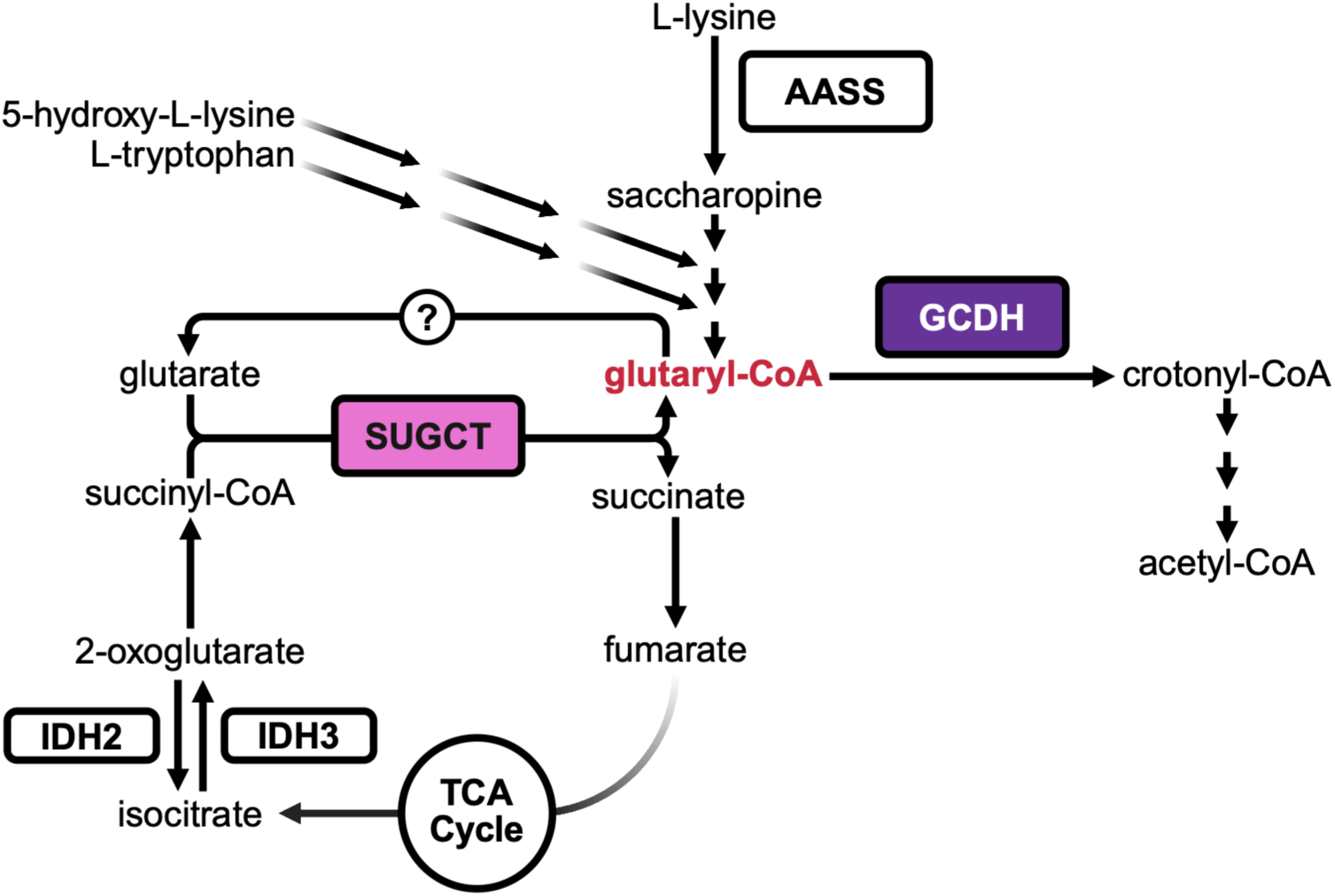
Simplified schema of lysine, hydroxylysine, and tryptophan catabolism.^3,4,7^. Lysine is degraded via two main catabolic pathways: The predominant mitochondrial saccharopine pathway and the enigmatic pipecolate pathway.^32,33^ The saccharopine pathway leads to the production of acetyl-CoA, with glutaryl-CoA as an intermediate; 5-hydroxy-L-lysine and tryptophan converge on this pathway upstream of glutaryl-CoA production.^4^ Because glutaryl-CoA is subject to spontaneous or enzyme-mediated hydrolysis to glutaric acid^34,35^, it must re-esterified to glutaryl-CoA by SUGCT. It is this regeneration that allows for the subsequent oxidative decarboxylation of glutaryl-CoA to crotonyl-CoA by GCDH.^36^

GA-1 (MIM 231670) is an autosomal recessive neurometabolic disorder caused by variants in the gene encoding glutaryl-CoA dehydrogenase (*GCDH*), which leads to accumulation of glutaryl-CoA, glutarylcarnitine (C5DC), and potentially neurotoxic organic acids such as 3-hydroxyglutaric and glutaric acid.^7–11^ The hallmark feature of untreated GA-1 is acute striatal injury, which leads to a complex movement disorder; other neurological manifestations may also develop in later-onset cases.^12–14^ Because immediate intervention is critical, GA-1 is included in newborn screening^15^, but some patients will still develop neurological damage. Therefore, GA-1 patients have an unmet medical need for an effective therapy, ideally one that would address the underlying metabolic defect responsible for the disease. Because SUGCT mediates the re-synthesis of glutaryl-CoA, inhibiting its catalytic activity could reduce the intracellular pool of glutaryl-CoA, thereby potentially mitigating downstream neurotoxicity without causing toxic systemic side effects. Therefore, we have proposed SUGCT as a novel therapeutic target in GA-1. Subsequently, we developed novel *in vitro* biochemical and cellular assays, performed a pilot high-throughput screen of a part of an FDA-approved drug library, and identified losartan carboxylic acid and valsartan as inhibitors of SUGCT with low micromolar IC_50_ values. Furthermore, we solved the crystal structures of SUGCT, both apo and with losartan carboxylic acid bound.^3^ However, neither losartan carboxylic acid nor valsartan successfully inhibited SUGCT in our cell models.

Herein, we address these challenges by developing an improved cell line assay for detecting SUGCT inhibition using a stable isotope-labeled glutarate (GA-d_4_). We also screened libraries of known bioactive compounds and identified additional SUGCT inhibitors for development as potential GA-1 therapeutics. Finally, we used the ethyl ester of valsartan as a prodrug to improve cell permeability and demonstrate the efficacy of the target. The new cellular assay and inhibitors provide a biochemical toolkit for screening small-molecule libraries to discover potent, specific SUGCT inhibitors for development as chemical probes or novel therapeutics for metabolic disorders such as GA-1.

## RESULTS

### Screening of the FDA-approved Drug Library

Utilizing our enzymatic SUGCT assay (Figure S1B) optimized in a 384-well format, we screened 469 compounds at a screening concentration of approximately 10 µM across two plates of our FDA-approved drug library for potential SUGCT inhibitors. Across both plates, we identified five statins as positive control hits, because the assay uses HMGCR as auxiliary enzyme. We also identified and purchased four hits as potential SUGCT inhibitors: Afatinib, meclofenamic acid, nafamostat mesylate, and panobinostat (Figure 2A).

**Figure 2:**
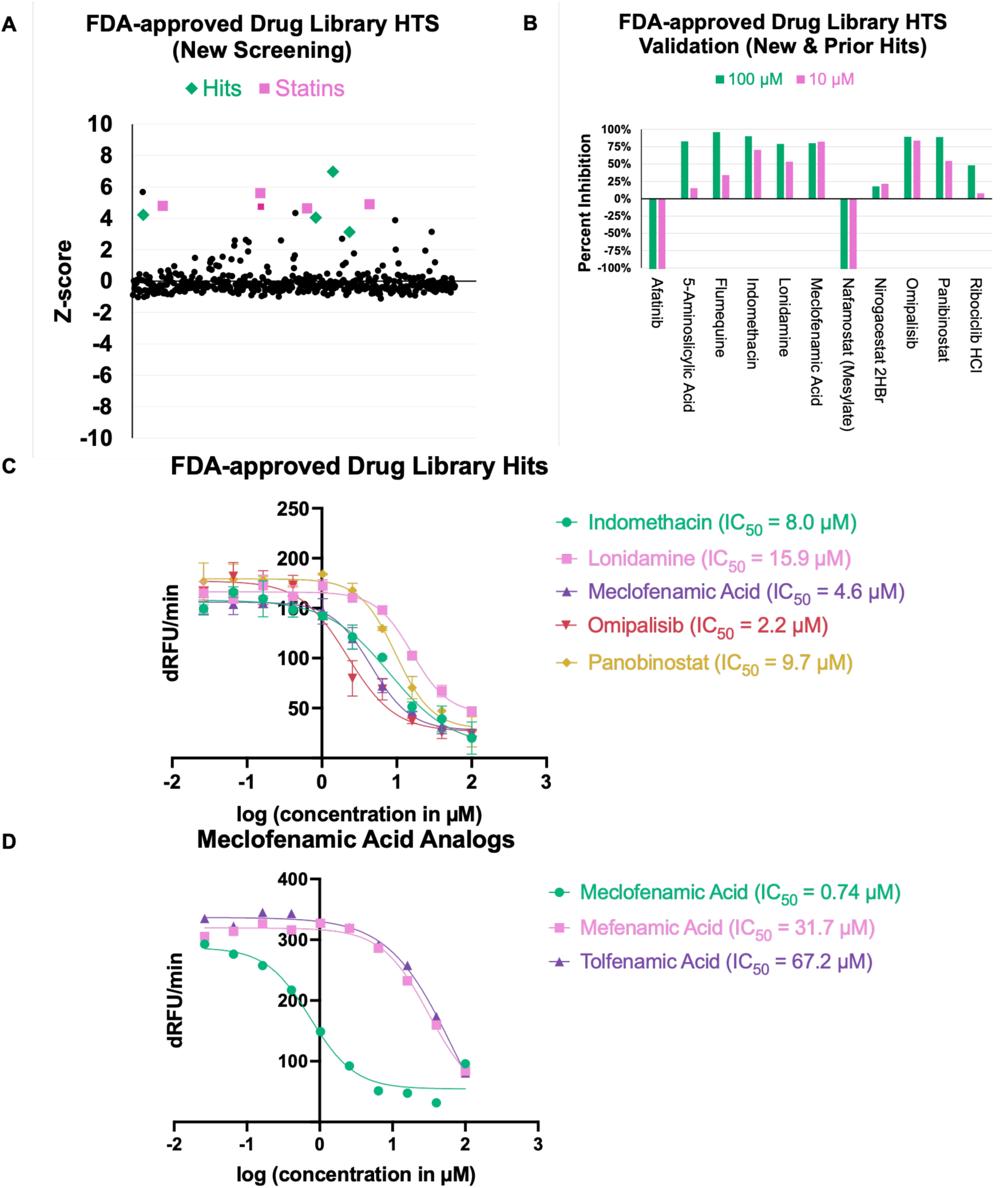
HTS & validation of FDA-approved drugs as potential SUGCT inhibitors. (A) Z-scores of screened compounds calculated individually utilizing the mean and standard deviation of all screening wells (except controls) excluding outliers as determined by compound fluorescence or quenching. Statins are marked by _▪_ in pink and hits are marked as • in green. (B) Re-purchase and validation of FDA-approved drugs. Each compound was measured in the SUGCT enzyme assay in duplicate at both 100 µM (green) and 10 µM (pink). (C) Dose-response curves for the top inhibitors from screening. (D) Dose-response curves for meclofenamic acid, mefenamic acid, and tolfenamic acid.

Upon purchasing these four hits for validation, we also purchased additional hits from our previously reported screening of FDA-approved drugs^3^: 5-aminosalicylic acid, flumequine, indomethacin, lonidamine, nirogacestat dihydrobromide, omipalisib, and ribociclib hydrochloride (Figure 2B). We immediately excluded afatinib and nafamostat (mesylate) from further testing because of outlier values at both 100 µM and 10 µM, most likely due to an assay artifact. Indomethacin, lonidamine, meclofenamic acid, omipalisib, and panobinostat all had greater than 50% inhibition at 10 µM, and were used to determine their IC_50_. These five compounds demonstrated SUGCT inhibition with IC_50_ values ranging from approximately 2 µM to 16 µM (Figure 2C, Table S1).

### Screening of the Bioactive Compound Library

In addition to the FDA-approved drug library, we screened 2,662 compounds between 8 plates of a bioactive compound library. From this library, we identified three statins as positive control hits and identified and purchased five additional hits for validation: Berberine hydrochloride, ertapenem sodium, GSK-J1, prulifloxacin, and UM 171 (Figure 3A). Because the IC_50_ of meclofenamic acid falls in the low micromolar range, we validated these compounds at 10 µM and 1 µM, rather than 100 µM and 10 µM (Figure 3B). Prulifloxacin was the only hit to inhibit SUGCT at ∼50% at 10 µM. Prulifloxacin (the prodrug of ulifloxacin) had an IC_50_ value of 2.6 µM, comparable to meclofenamic acid (Figure 3C, Table S1).

**Figure 3:**
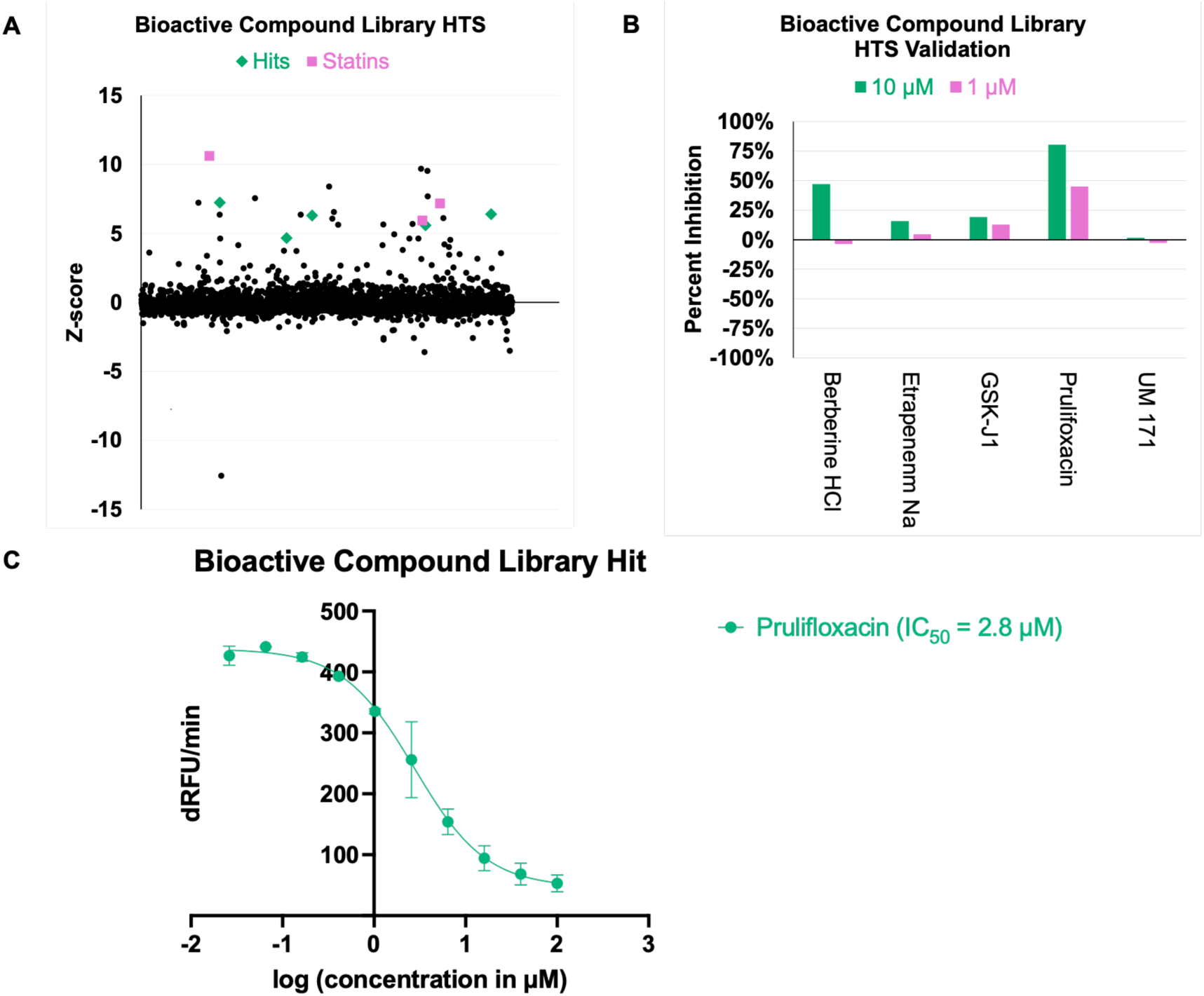
HTS & validation of bioactive compounds as potential SUGCT inhibitors. (A) Z-scores of screened compounds calculated individually utilizing the mean and standard deviation of all screening wells (except controls) excluding outliers as determined by compound fluorescence or quenching. Statins are marked by _▪_ in pink and hits are marked as • in green. (B) Re-purchase and validation of bioactive compounds. Each compound was measured in the SUGCT enzyme assay in duplicate at both 10 µM (green) and 1 µM (pink). (C) Dose-response curve for the top inhibitor from screening.

### Cellular Assay for SUGCT Activity Using Stable Isotope-Labeled Glutarate

Since endogenous *SUGCT* expression in HEK-293 cells is low, we had previously used the Flp-In system to generate HEK-293 cell lines stably overexpressing high levels of *SUGCT*. We then used CRISPR-Cas9 genome editing to create a *GCDH* KO in these *SUGCT*-overexpressing 293 cells, which caused a modest increase in C5DC accumulation. Unfortunately, we did not observe a decrease in C5DC upon treatment of *SUGCT*-overexpressing GA-1 HEK-293 cells with valsartan or its presumably more permeable ethyl ester prodrug. We had previously noted that *SUGCT*-overexpressing cell lines accumulate more C5DC upon addition of glutarate to the cell culture media. We reasoned that discriminating between glutaryl-CoA derived from endogenous lysine metabolism versus exogenous glutarate would improve the sensitivity of this assay. To test this hypothesis, we compared Flp-In-293 parental cells, *GCDH* KO, *SUGCT*-overexpressing cells, and *GCDH* KO cells with *SUGCT* overexpression (Figure 4A). These cells were incubated with regular DMEM or DMEM containing 5 mM glutarate (2,2,4,4-d_4_, 98%). In regular DMEM, *GCDH* KO increased C5DC accumulation, but there was no impact of *SUGCT* overexpression. The addition of glutarate-d_4_ to the media led to production of C5DC (M+4) proving that glutarate is re-esterified to glutaryl-CoA in HEK-293 cells. *GCDH* KO and *SUGCT* overexpression increased the accumulation of C5DC (M+4), but their combination had a synergistic effect, causing a very pronounced accumulation of C5DC (M+4).

**Figure 4:**
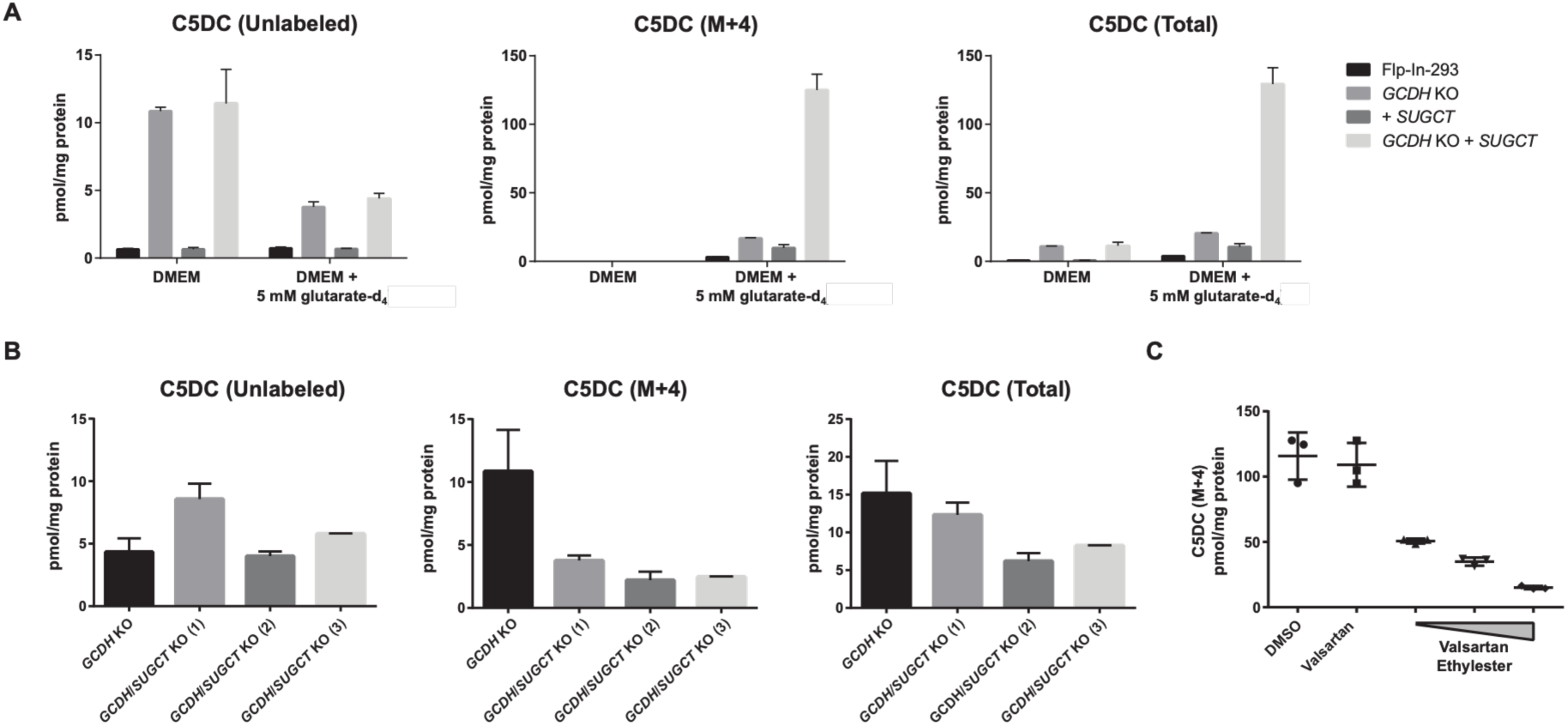
Development of cellular SUGCT activity assay. (A) Levels of unlabeled C5DC, C5DC (M+4) and the sum of both (total) in cell pellets after 24 hours of incubation in DMEM or DMEM supplemented with 5 mM glutarate-d_4_. Four different cell lines were compared; parental Flp-In-293 cells, *GCDH* KO cells, *SUGCT* overexpressing cells, and *SUGCT* overexpressing cells with *GCDH* KO. (B) C5DC levels in cell pellets after 24 hours of incubation in DMEM supplemented with 5 mM glutarate-d_4_. Four different cell lines were compared; parental *GCDH* KO cells and three different *GCDH*/*SUGCT* KO cell lines. (C) C5DC levels in pellets of *SUGCT* overexpressing cells with *GCDH* KO after 24 hours of incubation in DMEM supplemented with 5 mM glutarate-d_4_. The concentration of valsartan was 50 µM, the concentrations for valsartan ethyl ester were 8, 20 and 50 µM. For all graphs, mean and SD are displayed. Each condition was performed in triplicate.

Even Flp-In-293 parental cells and *GCDH* KO accumulated some C5DC (M+4) upon loading with glutarate-d_4_, suggesting that there is endogenous *SUGCT* expression in HEK-293 cells. To provide additional proof, we used three previously generated *GCDH*/*SUGCT* double KO cell lines (Figure 4B). The homozygous mutations in *SUGCT* are: (1) a 2 bp insertion (p.V189Lfs*5), (2) a 51 bp deletion (removing amino acids 171-187), and (3) a 26 bp deletion (p.V185Afs*6). We compared C5DC (M+4) accumulation after loading with glutarate-d_4_ to the parental *GCDH* KO clone. C5DC (M+4) levels decreased upon *SUGCT* KO, proving that endogenous *SUGCT* can be a source of C5DC in HEK-293 cells. It is noteworthy that even in *GCDH*/*SUGCT* double KO cells, C5DC (M+4) formation remains detectable suggesting that there are other enzymes able to re-esterify glutarate.

To determine if these newly established cell line conditions with 5 mM glutarate-d_4_ can be used to detect inhibition of SUGCT activity, we used first compared the effects of valsartan and its ethyl ester on C5DC (M+4) formation in *GCDH* KO cells that overexpress *SUGCT* (Figure 4C). At 50 µM, valsartan was inactive, whereas valsartan ethyl ester decreased C5DC (M+4) dose-dependently, with 8 µM already producing more than a 50% decrease. These data demonstrate that the adapted cell line model enables the discovery of cell-permeable SUGCT inhibitors.

### In-cell Inhibition of SUGCT by Meclofenamic Acid

Having shown that our cell assay using stable isotope-labeled glutarate can detect inhibition of SUGCT activity, we then tested each of the hits used in IC_50_ experiments for their cellular efficacy. At 50 µM, omipalisib and panobinostat were deemed toxic due to changes in cell morphology, whereas lonidamine did not reduce C5DC (M+4) accumulation; however, meclofenamic acid lowered C5DC (M+4) levels (Figure 4A). Thus, we tested meclofenamic acid at varying concentrations against a positive control, valsartan ethyl ester (Figure 4B). In a similar manner to valsartan ethyl ester, meclofenamic acid reduced C5DC (M+4) dose-dependently, yet toxicity was observed at 100 µM. Prulifloxacin did not reduce C5DC (M+4) levels at 50 µM, and its active metabolite, ulifloxacin, increased C5DC (M+4) levels (Figure 4C).

### Further Characterization of the SUGCT:Meclofenamic Acid Complex

Because meclofenamic acid was able to inhibit SUGCT in our cell assay (Figure 4A), we proceeded with its further characterization. We tested four of its analogs alongside meclofenamic acid in our enzyme assay: Fenamic acid, flufenamic acid, mefenamic acid, and tolfenamic acid. These compounds are all fenamates, which are derived from the fenamic acid core structure, which are a subgroup of clinically prescribed NSAIDs that are also studied for off-label applications in various diseases, including cancer and Alzheimer’s.^16^ Fenamic acid and flufenamic acid did not inhibit SUGCT, while mefenamic acid and tolfenamic acid showed 42.5-fold and 90-fold reductions in potency, respectively, compared to meclofenamic acid (Figure 1D).

To further characterize meclofenamic acid binding to SUGCT, we attempted to crystallize the complex as under previous conditions^3^, using both wild-type SUGCT and the Q255A/K256A mutant, but we were unable to obtain diffracting crystals. To investigate the structural properties of meclofenamic acid binding to SUGCT, we performed a Boltz-2 binding prediction in ChimeraX for meclofenamic acid and our previous apo SUGCT structure (PDB: 9BR6^3^, Table S2).

SUGCT forms its active site via the interface of two monomers^3^, and our model predicted meclofenamic acid binding a pocket within this region (Figures 5A, 5B), with two predicted hydrogen bonds to the backbone nitrogen of G162 (3.2 Å) and G174 (3.1 Å) (Figure 5C, S3).

**Figure 5:**
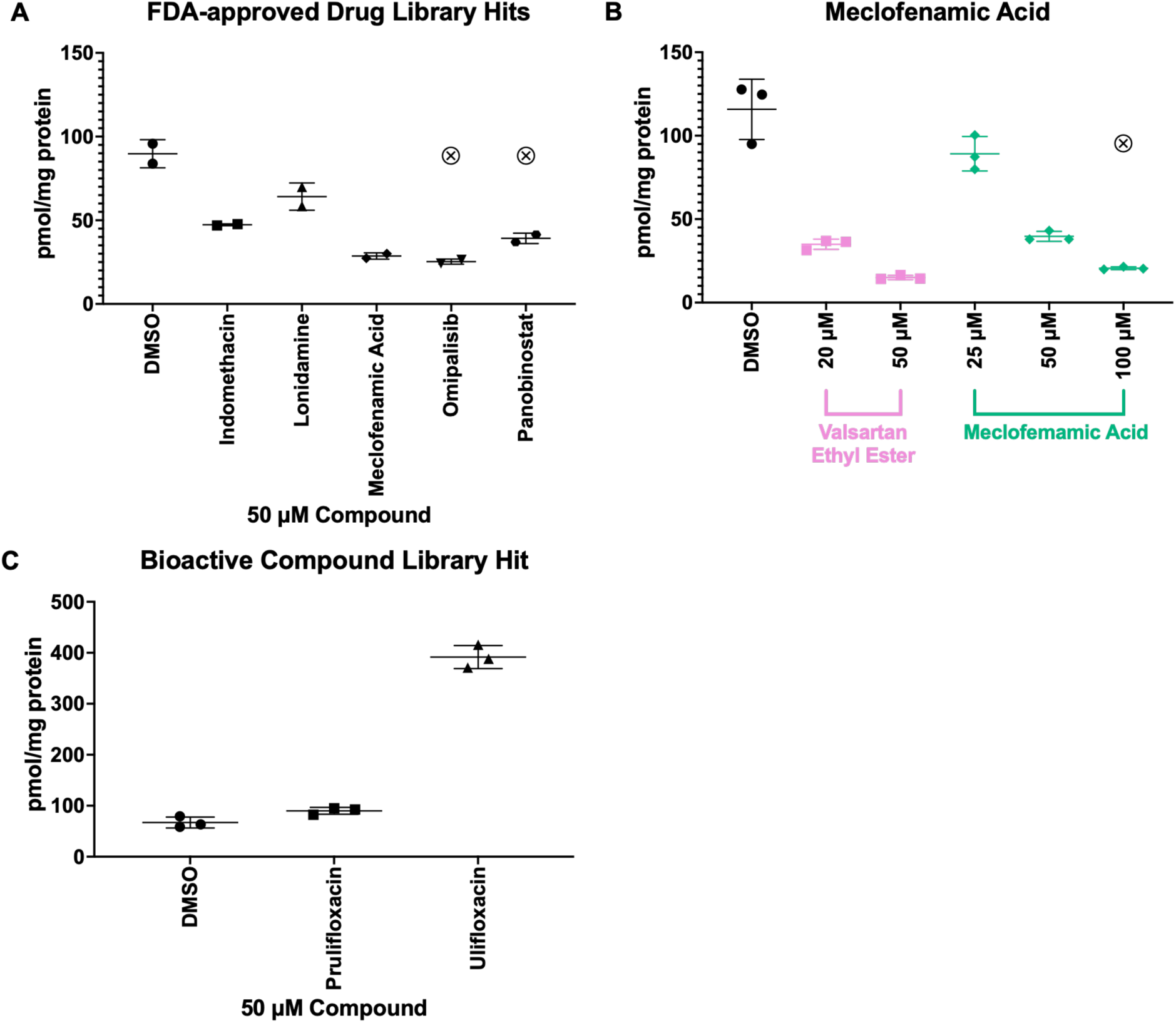
Cellular activity of hit compounds. (A) Hits from the FDA-approved drug library tested in the cell assay at 50 µM. Toxicity noted by ⊗. (B) Varying concentrations of meclofenamic acid (green) tested in the cell assay against valsartan ethyl ester (pink) as a positive control. Toxicity noted by ⊗. (C) Hits from the bioactive compound library tested at 50 µM.

**Figure 6:**
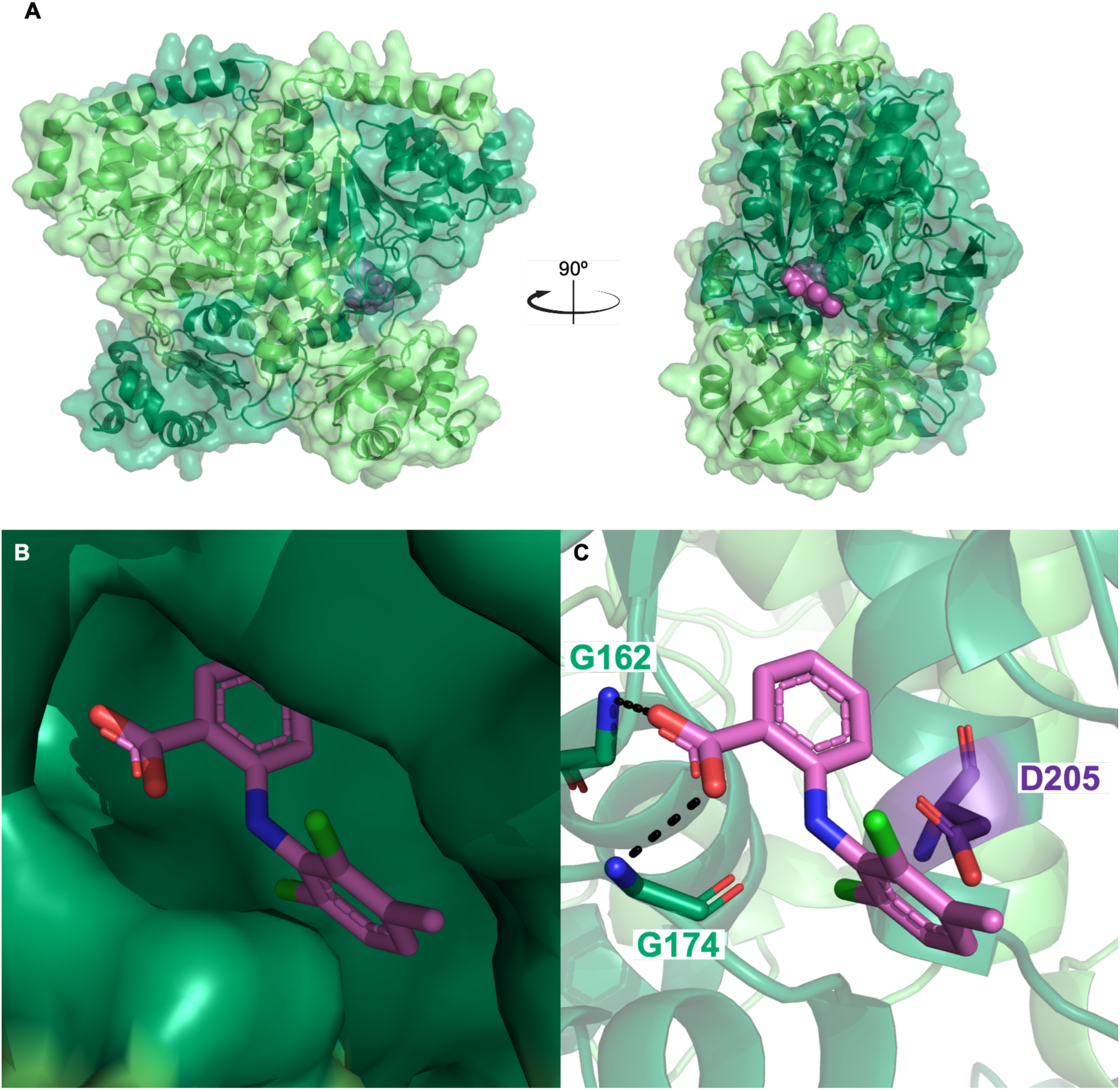
Boltz-2 prediction of the SUGCT:Meclofenamic acid complex. (A) Ribbon and surface model of SUGCT (chain A in dark green, chain B in light green) with meclofenamic acid bound (pink spheres). (C) Close-up of a surface model of the interaction of meclofenamic acid (pink carbon backbone) and SUGCT. (B) Close-up of the interaction of meclofenamic acid with G162 and G174 of SUGCT, with catalytic residue D205 (purple backbone) highlighted.

Meclofenamic acid is predicted to bind closely to the highly conserved catalytic residue D205^3^ (Figure 5C, S3). To explore this further, we aligned the highly conserved homolog Formyl-coenzyme A transferase from *Brucella melitensis*, in complex with both succinate (PDB: 9ZAV^17^) and CoA (PDB: 10UH^18^), from experimental crystal structures in the PDB (Figure S4, S5). Notably, our model’s ligand directly clashes with the bound CoA in 9ZAV and the bound succinate in 10UH.

## DISCUSSION

Utilizing our previously developed SUGCT enzymatic assay, we screened 3,131 compounds for inhibition and observed a hit rate of ∼ 0.28%. Meclofenamic acid inhibited SUGCT both enzymatically and in cells, but was toxic at higher concentrations. Interestingly, meclofenamic acid has been reported to disrupt mitochondrial respiration *in vivo*^19^, suggesting that this phenomenon, or a similar one, may be occurring in HEK-293 cells treated with meclofenamic acid and leading to downstream toxicity. We suggest that an ethyl ester of meclofenamic acid may be more permeable, which could reduce the observed toxicity.

Furthermore, the Boltz-2 model of the SUGCT:Meclofenamic Acid complex yielded useful insights. While further biophysical insights are needed, the promiscuity of the active site of SUGCT, the Boltz-2 prediction, as well as the prediction’s alignment with experimental structures of homologs bound to their endogenous substrates, indicate a likelihood of competitive inhibition. Conversely to our previously reported inhibitor, losartan carboxylic acid, meclofenamic acid is predicted to bind directly to SUGCT, rather than rely on coordination of salt ions in the active site. While the IC_50_ of losartan carboxylic acid was dependent on the presence of potassium and phosphate ions *in vitro*, no such change was observed for meclofenamic acid (Figure S6), which in turn supports the Boltz-2 prediction.

Prulifloxacin was not active in our cell assay, whereas ulifloxacin, its active metabolite, paradoxically increased C5DC levels. Perhaps the discordance between these results is explained by limited expression of paraoxonase(s) in HEK-293 cells, thereby resulting in inadequate hydrolysis of prulifloxacin to ulifloxacin.^20^ Moreover, ulifloxacin, which biochemically inhibited SUGCT with an IC_50_ of approximately 5.2 µM (Figure S7), increased C5DC (M+4) levels. While evidently not a SUGCT activator, ulifloxacin may be an undocumented activator of a transporter that our cell assay utilizes for uptake of glutarate present in the cell media (Figure S8). Glutarate is a transported substrate of both the sodium-dependent dicarboxylate transporter NaDC3^23^ (which is essential for cellular uptake of TCA cycle intermediates, including succinate^24^) and the mitochondrial 2-oxodicarboxylate carrier SLC25A21^25^. Additionally, ulifloxacin may activate a carnitine acyl transferase (CAT), which is necessary for the conversion of glutaryl-CoA to C5DC, which can then cross membranes.^26,27^ Alternatively, it display other off-target specificity causing disruptions in mitochondrial function. Recent work has identified fluoroquinolones, the class of antibiotics to which ulifloxacin belongs, as prominent disruptors of mitochondrial metabolism in HEK-293 cells^28^, which may, in turn, play a role in glutaryl-CoA and carnitine homeostasis.

Together, these findings have furthered the development of cell-permeable inhibitors for SUGCT. Valsartan ethyl ester and meclofenamic acid are cell-permeable inhibitors of SUGCT that are able to reduce C5DC levels in cells and can be useful tool compounds to further study the role of SUGCT in GA-1. Further optimization of these compounds and further exploration of them as scaffolds may yield inhibitors that demonstrate both permeability and high cellular potency, which may be used as chemical probes for SUGCT activity and/or further developed into potential GA-1 therapeutics in the future.

## METHODS

### Materials

The FDA-approved drug library and bioactive compound library were obtained from APExBIO. All individual compounds were purchased from MedChemExpress.

### Purification of Recombinant SUGCT

To produce recombinant SUGCT protein for enzymatic assays, we utilized an N-terminal His tag in a pET47 vector, as we had done in the past. We expressed this His-SUGCT in pGro7 cells, inoculating 2 L of TB with 5 mL overnight culture and 40 μg/ml kanamycin. Then, we grew the cells to an OD_600_ of approximately 1.5, cooled to 16°C, and induced overnight with 400 _μM_ IPTG and 20 mL of 40% arabinose. The next day, we resuspended the pellet in TBS and 10% glycerol, along with PMSF and lysozyme, and then lysed it using a sonicator (Qsonica). After pelleting cellular debris, we loaded the lysate onto a column containing HisPur Ni-NTA Resin (Thermo Scientific) for immobilized metal affinity chromatography (IMAC). The column was equilibrated with 20 mM Tris pH 8.0, 250 mM NaCl, and 40 mM imidazole, then washed with 30 mL of 20 mM Tris 8.0, 250 mM NaCl, and 50 mM imidazole. The His-tagged protein was eluted with 20 mM Tris pH 8.0, 150 mM NaCl, 250 mM imidazole, and 10% glycerol. After elution, we added HRV3C protease and THP overnight. The following day, we purified the protein on a Superdex200 Increase size-exclusion column, concentrated, and flash-froze the protein with 10% glycerol for assays. All samples were analyzed by SDS-PAGE.

### Purification of Recombinant HMG-CoA Reductase (HMGCR)

The catalytic portion of human HMG-CoA reductase (HMGCR) was expressed with an N-terminal His-tag as described.^3^ Here we used Lemo cells (NEB), inoculating 2 L of LB with 5 mL overnight culture and 40 μg/ml kanamycin. Cells were then grown to an OD_600_ of approximately 1.0, cooled to 24°C, and induced overnight with 400 _μM_ IPTG. The following day, we purified by IMAC in the same manner as described for SUGCT. After elution, we purified the protein on a Superdex200 Increase size-exclusion column, concentrated, and froze the protein for assays with 10% glycerol. All samples were analyzed by SDS-PAGE.

### SUGCT Enzyme Assay

For high-throughput screening, validation, and IC_50_ experiments, we used the conditions described.^3^ Enzymes were added at differing concentrations based on their measured activity in a test assay following purification. Percent inhibition was calculated by subtracting

### Cell Assay

*SUGCT* was targeted using CRISPR-Cas9 genome editing as described.^29^ For this, a pCas9-GFP plasmid with a *SUGCT* guide sequence targeting exon 7 (Fw: 5’-CACCGCATCAGACCAGAAACAGCCG-3’ and Rev: 5’-AAACCGGCTGTTTCTGGTCTGATGC-3’) was transfected into a previously validated *GCDH* KO HEK-293 cell line using Lipofectamine 2000. After 48 h, transfected cells were collected and GFP positive single cells were sorted into a well of a 96 well plate with 50% conditioned DMEM (DMEM collected from the same cell line when cells were 60–90% confluent mixed 1:1 with fresh DMEM supplemented with Pen/Strep and 20% FBS). After 12–15 days of incubation at 37°C and 5% CO_2_, clones appeared in several wells. Clones were first transferred to 24-well plates and after reaching 70–90% confluence were transferred to T25 flasks. Cell pellets were then collected for confirmation of *SUGCT* KO through DNA amplification and Sanger sequencing using primers Fw: 5’-CTGCCTTCTTCAAGGAATGGT-3’ and Rev: 5’-AAGCATTCACTTTCCCTGGT-3’.

Glutaric acid (2,2,4,4-D4, 98%) was purchased from Cambridge Isotope Laboratories, Inc (Andover, MA). The labeled glutaric acid was dissolved in water and neutralized with KOH. The resulting 334 mM stock solution of labeled glutarate was diluted to 5 mM in DMEM media with 400 µM L-carnitine. After 24 hours incubation with these media, the intracellular metabolites were immediately extracted by adding 1 mL ice-cold 5:3:2 extraction solvent (methanol:acetonitrile:water) directly to the plate with cells and incubated overnight at −80°C to aid protein precipitation. The following day, the cells were scraped, and the extracts centrifuged at 20,000×g for 20 min at 4°C. Carnitine/Acylcarnitine Standards Mix B (NSK-B-1, Cambridge Isotope Laboratories) were added, and samples were analyzed by mass spectrometry as described.^29^

### Boltz-2 Prediction

Boltz-2 prediction was made using the sequence of the previously determined apo structure of SUGCT (PDB: 9BR6) with meclofenamic acid.^3,30^ The predicted complex was analyzed using UCSF ChimeraX.^31^ Alignments were performed of the predicted complex against experimentally determined crystal structures of formyl-coenzyme A transferase from *Brucella melitensis* (PDB: 9ZAV^17^ and PDB: 10UH^18^).

## Supporting information

All supplemental tables and figures

## ACKNOWLEDGMENTS

We thank Anna Liu, Samantha Pabua and Kedar Tandukar for their excellent technical assistance with the clinical biochemical analyses.

## REFERENCES

1. Marlaire, S.; Van Schaftingen, E.; Veiga-da-Cunha, M. C7orf10 Encodes Succinate-Hydroxymethylglutarate CoA-Transferase, the Enzyme That Converts Glutarate to Glutaryl-CoA. Journal of Inherited Metabolic Disease 2014, 37, 13–19. 10.1007/s10545-013-9632-0.

2. Sherman, E. A.; Strauss, K. A.; Tortorelli, S.; Bennett, M. J.; Knerr, I.; Morton, D. H.; Puffenberger, E. G. Genetic Mapping of Glutaric Aciduria, Type 3, to Chromosome 7 and Identification of Mutations in C7orf10. American Journal of Human Genetics 2008, 83, 604–609. 10.1016/j.ajhg.2008.09.018.

3. Wu, R.; Khamrui, S.; Dodatko, T.; Leandro, J.; Sabovic, A.; Violante, S.; Cross, J. R.; Marsan, E.; Kumar, K.; DeVita, R. J.; Lazarus, M. B.; Houten, S. M. Characterization, Structure, and Inhibition of the Human Succinyl-CoA:Glutarate-CoA Transferase, a Putative Genetic Modifier of Glutaric Aciduria Type 1. ACS Chemical Biology 2024, 19, 1544–1553. 10.1021/acschembio.4c00204.

4. Hagen, J.; te Brinke, H.; Wanders, R. J. A.; Knegt, A. C.; Oussoren, E.; Hoogeboom, A. J.; Ruijter, G. J. G.; Becker, D.; Schwab, K. O.; Franke, I.; Duran, M.; Waterham, H. R.; Sass, J. O.; Houten, S. M. Genetic Basis of Alpha-Aminoadipic and Alpha-Ketoadipic Aciduria. Journal of Inherited Metabolic Disease 2015, 38, 873–879. 10.1007/s10545-015-9841-9.

5. Linster, C. L.; Van Schaftingen, E.; Hanson, A. D. Metabolite Damage and Its Repair or Pre-Emption. Nature Chemical Biology 2013, 9, 72–80. 10.1038/nchembio.1141.

6. Knerr, I.; Zschocke, J.; Trautmann, U.; Dorland, L.; de Koning, T. J.; Müller, P.; Christensen, E.; Trefz, F. K.; Wündisch, G. F.; Rascher, W.; Hoffmann, G. F. Glutaric Aciduria Type III: A Distinctive Non-Disease? Journal of Inherited Metabolic Disease 2002, 25, 483–490. 10.1023/A:1021207419125.

7. Leandro, J.; Bender, A.; Dodatko, T.; Argmann, C.; Yu, C.; Houten, S. M. Glutaric Aciduria Type 3 Is a Naturally Occurring Biochemical Trait in Inbred Mice of 129 Substrains. Molecular Genetics and Metabolism 2021, 132, 139–145. 10.1016/j.ymgme.2021.01.004.

8. Kölker, S.; Koeller, D. M.; Okun, J. G.; Hoffmann, G. F. Pathomechanisms of Neurodegeneration in Glutaryl-CoA Dehydrogenase Deficiency. Annals of Neurology 2004, 55, 7–12. 10.1002/ana.10784.

9. Kölker, S.; Ahlemeyer, B.; Krieglstein, J.; Hoffmann, G. F. Maturation-Dependent Neurotoxicity of 3-Hydroxyglutaric and Glutaric Acids in Vitro: A New Pathophysiologic Approach to Glutaryl-CoA Dehydrogenase Deficiency. Pediatric Research 2000, 47, 495–503. 10.1203/00006450-200004000-00014.

10. Goodman, S. I.; Kratz, L. E.; DiGiulio, K. A.; Biery, B. J.; Goodman, K. E.; Isaya, G.; Frerman, F. E. Cloning of Glutaryl-CoA Dehydrogenase cDNA, and Expression of Wild Type and Mutant Enzymes in Escherichia coli. Human Molecular Genetics 1995, 4, 1493–1498. 10.1093/hmg/4.9.1493.

11. Goodman, S. I.; Stein, D. E.; Schlesinger, S.; Christensen, E.; Schwartz, M.; Greenberg, C. R.; Elpeleg, O. N. Glutaryl-CoA Dehydrogenase Mutations in Glutaric Acidemia (Type I): Review and Report of Thirty Novel Mutations. Human Mutation 1998, 12, 141–144

12. Strauss, K. A.; Puffenberger, E. G.; Robinson, D. L.; Morton, D. H. Type I Glutaric Aciduria, Part 1: Natural History of 77 Patients. American Journal of Medical Genetics Part C: Seminars in Medical Genetics 2003, 121C, 38–52. 10.1002/ajmg.c.20007.

13. Kölker, S.; Christensen, E.; Leonard, J. V.; Greenberg, C. R.; Boneh, A.; Burlina, A. B.; Burlina, A. P.; Dixon, M.; Duran, M.; García Cazorla, A.; Goodman, S. I.; Koeller, D. M.; Kyllerman, M.; Mühlhausen, C.; Müller, E.; Okun, J. G.; Wilcken, B.; Hoffmann, G. F.; Burgard, P. Diagnosis and Management of Glutaric Aciduria Type I—Revised Recommendations. Journal of Inherited Metabolic Disease 2011, 34, 677–694. 10.1007/s10545-011-9289-5.

14. Boy, N.; Garbade, S. F.; Heringer, J.; Seitz, A.; Kölker, S.; Harting, I. Patterns, Evolution, and Severity of Striatal Injury in Insidious-vs Acute-Onset Glutaric Aciduria Type 1. Journal of Inherited Metabolic Disease 2019, 42, 117–127. 10.1002/jimd.12033.

15. Boy, N.; Mengler, K.; Thimm, E.; Schiergens, K. A.; Marquardt, T.; Weinhold, N.; Marquardt, I.; Das, A. M.; Freisinger, P.; Grünert, S. C.; Vossbeck, J.; Steinfeld, R.; Baumgartner, M. R.; Beblo, S.; Dieckmann, A.; Näke, A.; Lindner, M.; Heringer, J.; Hoffmann, G. F.; Mühlhausen, C.; Maier, E. M.; Ensenauer, R.; Garbade, S. F.; Kölker, S. Newborn Screening: A Disease-Changing Intervention for Glutaric Aciduria Type 1. Annals of Neurology 2018, 83, 970–979. 10.1002/ana.25233.

16. Hill, J.; Zawia, N. H. Fenamates as Potential Therapeutics for Neurodegenerative Disorders. Cells 2021, 10, 702. 10.3390/cells10030702.

17. Seattle Structural Genomics Center for Infectious Disease (SSGCID) Crystal Structure of Formyl-Coenzyme A Transferase from *Brucella melitensis* in Complex with Succinate. Protein Data Bank 2025, PDB ID 9ZAV. 10.2210/pdb9ZAV/pdb.

18. Seattle Structural Genomics Center for Infectious Disease (SSGCID) Crystal Structure of Formyl-Coenzyme A Transferase from *Brucella melitensis* in Complex with CoA. Protein Data Bank 2026, PDB ID 10UH. 10.2210/pdb10UH/pdb.

19. Rodríguez-Enríquez, S.; Robledo-Cadena, D. X.; Pacheco-Velázquez, S. C.; Vargas-Navarro, J. L.; Padilla-Flores, J. A.; Kaambre, T.; Moreno-Sánchez, R. Repurposing Auranofin and Meclofenamic Acid as Energy-Metabolism Inhibitors and Anti-Cancer Drugs. PLOS ONE 2024, 19 (9), e0309331. 10.1371/journal.pone.0309331

20. Tougou, K.; Nakamura, A.; Watanabe, S.; Okuyama, Y.; Morino, A. Paraoxonase Has a Major Role in the Hydrolysis of Prulifloxacin (NM441), a Prodrug of a New Antibacterial Agent. Drug Metabolism and Disposition 1998, 26 (4), 355–359.

21. Sood, R.; Flint-Ashtamker, G.; Borenstein, D.; Barki-Harrington, L. Upregulation of Prostaglandin Receptor EP1 Expression Involves Its Association with Cyclooxygenase-2. PLOS ONE 2014, 9 (3), e91018. 10.1371/journal.pone.0091018

22. Matsunobu, T.; Okuno, T.; Yokoyama, C.; Yokomizo, T. Thromboxane A Synthase-Independent Production of 12-Hydroxyheptadecatrienoic Acid, a BLT2 Ligand. Journal of Lipid Research 2013, 54 (11), 2979–2987. 10.1194/jlr.M037754

23. Stellmer, F.; Keyser, B.; Burckhardt, B. C.; Koepsell, H.; Streichert, T.; Glatzel, M.; Jabs, S.; Thiem, J.; Herdering, W.; Koeller, D. M.; Goodman, S. I.; Lukacs, Z.; Ullrich, K.; Burckhardt, G.; Braulke, T.; Mühlhausen, C. 3-Hydroxyglutaric Acid Is Transported via the Sodium-Dependent Dicarboxylate Transporter NaDC3. Journal of Molecular Medicine 2007, 85, 763–770. 10.1007/s00109-007-0174-5.

24. Li, P.; Chen, B.; Dong, M.; Luo, J.; Lin, N.; Li, Y. SLC13 Sodium-Carboxylate Transporters: Function, Regulation and Pathophysiological Implications in Human Disease. Biochemical Pharmacology 2026, 246, 117744. 10.1016/j.bcp.2026.117744.

25. Fiermonte, G.; Dolce, V.; Palmieri, L.; Ventura, M.; Runswick, M. J.; Palmieri, F.; Walker, J. E. Identification of the Human Mitochondrial Oxodicarboxylate Carrier: Bacterial Expression, Reconstitution, Functional Characterization, Tissue Distribution, and Chromosomal Location. Journal of Biological Chemistry 2001, 276, 8225–8230. 10.1074/jbc.M009607200.

26. Jennings, E. A.; Abi-Rached, Z. H.; Jones, D. E.; Ryan, R. O. 3-Methylglutarylcarnitine: A Biomarker of Mitochondrial Dysfunction. Clinica Chimica Acta 2023, 551, 117629. 10.1016/j.cca.2023.117629.

27. Richert, A. C.; Zhang, Y.; Bharathi, S. S.; Hernandez, A.; Dodatko, T.; Bons, J.; Roundtree, K. E.; Stauffer, B.; Yu, C.; Schilling, B.; Houten, S. M.; Goetzman, E. S. Odd-Chain Dicarboxylic Acid Feeding Produces a Glutaric Aciduria Type 1-Like Metabolic Signature in Mice. The FASEB Journal 2025, 39, e71215. 10.1096/fj.202502381R.

28. Reinhardt, T.;, et al. Chemical Proteomics Reveals Human Off-Targets of Fluoroquinolone Induced Mitochondrial Toxicity. Angewandte Chemie International Edition 2025, 64 (18), e202421424. 10.1002/anie.202421424.

29. Leandro, J.; Dodatko, T.; Aten, J.; Nemeria, N. S.; Zhang, X.; Jordan, F.; Hendrickson, R. C.; Sanchez, R.; Yu, C.; DeVita, R. J.; Houten, S. M. DHTKD1 and OGDH Display Substrate Overlap in Cultured Cells and Form a Hybrid 2-Oxo Acid Dehydrogenase Complex *In Vivo*. Human Molecular Genetics 2020, 29 (7), 1168–117.

30. Passaro, S.; Corso, G.; Wohlwend, J.; Reveiz, M.; Thaler, S.; Somnath, V. R.; Getz, N.; Portnoi, T.; Roy, J.; Stark, H.; Kwabi-Addo, D.; Beaini, D.; Jaakkola, T.; Barzilay, R. Boltz-2: Towards Accurate and Efficient Binding Affinity Prediction. bioRxiv 2025, 2025.06.14.659707. 10.1101/2025.06.14.659707.

31. Pettersen, E. F.; Goddard, T. D.; Huang, C. C.; Meng, E. C.; Couch, G. S.; Croll, T. I.; Morris, J. H.; Ferrin, T. E. UCSF ChimeraX: Structure Visualization for Researchers, Educators, and Developers. Protein Science 2021, 30, 70–82. 10.1002/pro.3943.

32. Hallen, A.; Jamie, J. F.; Cooper, A. J. L. Lysine Metabolism in Mammalian Brain: An Update on the Importance of Recent Discoveries. Amino Acids 2013, 45, 1249–1272. 10.1007/s00726-013-1590-1.

33. Pena, I. A.; Marques, L. A.; Laranjeira, A. B. A.; Yunes, J. A.; Eberlin, M. N.; MacKenzie, A.; Arruda, P. Mouse Lysine Catabolism to Aminoadipate Occurs Primarily through the Saccharopine Pathway; Implications for Pyridoxine Dependent Epilepsy (PDE). Biochimica et Biophysica Acta - Molecular Basis of Disease 2017, 1863, 121–128. 10.1016/j.bbadis.2016.09.006.

34. Westin, M. A. K.; Hunt, M. C.; Alexson, S. E. H. The Identification of a Succinyl-CoA Thioesterase Suggests a Novel Pathway for Succinate Production in Peroxisomes. Journal of Biological Chemistry 2005, 280, 38125–38132. 10.1074/jbc.M508479200.

35. Wagner, G. R.; Bhatt, D. P.; O’Connell, T. M.; Thompson, J. W.; Dubois, L. G.; Backos, D. S.; Yang, H.; Mitchell, G. A.; Ilkayeva, O. R.; Stevens, R. D.; Grimsrud, P. A.; Hirschey, M. D. A Class of Reactive Acyl-CoA Species Reveals the Non-Enzymatic Origins of Protein Acylation. Cell Metabolism 2017, 25, 823–837.e8. 10.1016/j.cmet.2017.03.006.

36. Fu, Z.; Wang, M.; Paschke, R.; Rao, K. S.; Frerman, F. E.; Kim, J.-J. P. Crystal Structures of Human Glutaryl-CoA Dehydrogenase with and without an Alternate Substrate: Structural Bases of Dehydrogenation and Decarboxylation Reactions. Biochemistry 2004, 43, 9674–9684. 10.1021/bi049290c.

37. Notredame, C.; Higgins, D. G.; Heringa, J. T-Coffee: A Novel Method for Fast and Accurate Multiple Sequence Alignment. Journal of Molecular Biology 2000, 302, 205–217. 10.1006/jmbi.2000.4042.

