## Supplementary material for "Discovery of cell-permeable inhibitors for human succinyl-CoA:glutarate-CoA transferase": All supplemental tables and figures

**Contents**:

- **SUPPLEMENTARY FIGURES & TABLES**

### SUPPLEMENTARY FIGURES & TABLES:

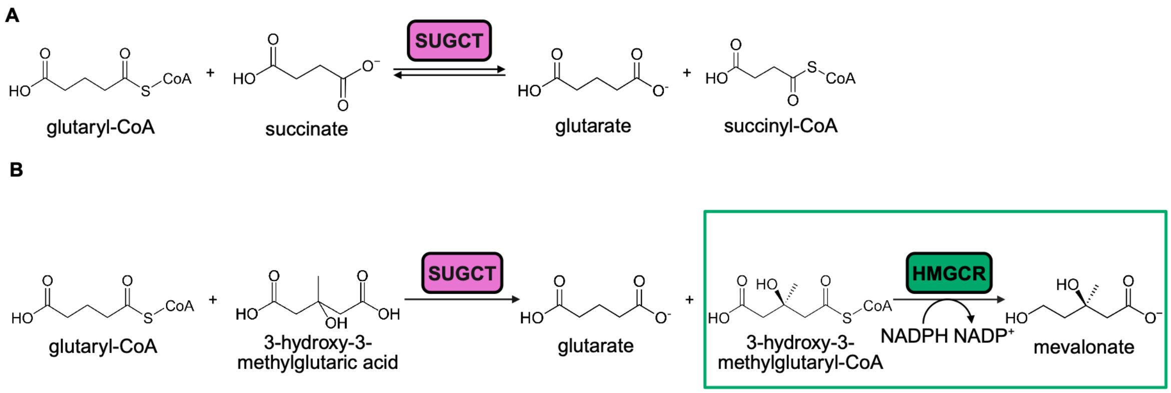


**Figure S1: SUGCT biochemistry.** (A) Physiological SUGCT reaction.[1,3] SUGCT mediates the reversible transfer of Coenzyme A between glutaryl-CoA and free succinate. (B) Schema of SUGCT enzyme assay.[1,3] SUGCT can utilize different carboxylic acids as substrates; here, SUGCT transfers Coenzyme A from glutaryl-CoA to 3-hydroxy-3-methyglutaric acid to form free glutarate and 3-hydroxy-3-methylglutaryl-CoA. This reaction is coupled with HMGCR, which, in turn, reduces 3-hydroxy-3-methylglutaryl-CoA to mevalonate, consuming two NADPH in the process. This NADPH consumption is measured fluorometrically, with excitation at 340 nm and emission at 460 nm, after addition of 3-hydroxy-3-methyglutaric acid to start the first reaction.

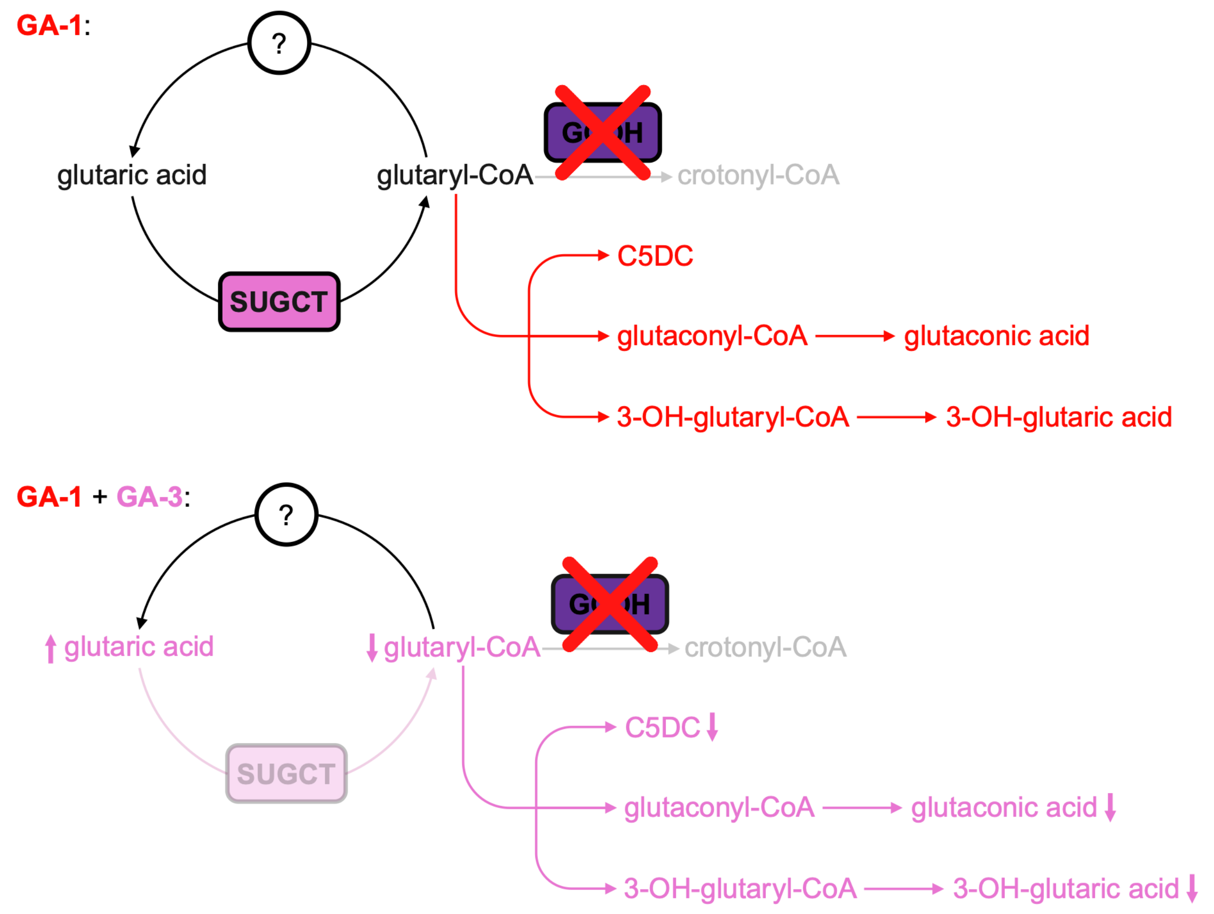


**Figure S2: Glutaric aciduria biochemical phenotypes as observed in mice.**[7] Glutaconic and 3-hydroxyglutaric acid are neurotoxic.[7,9]

**Table S1: IC$_{50}$ values for SUGCT inhibitors.**

| Compound Name | Compound Structure | IC$_{50}$ (μM) |
|---|---|---|
| Indomethacin | 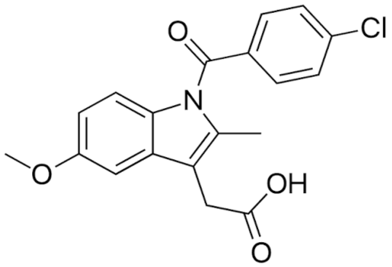 | 8.0 |
| Lonidamine | 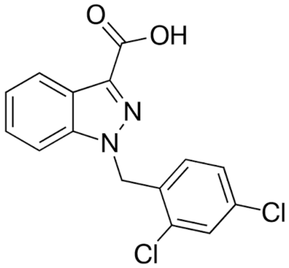 | 15.9 |
| Meclofenamic Acid | 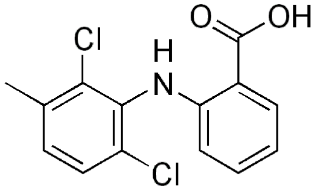 | 4.6 |
| Omipalisib | 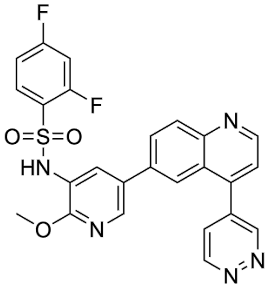 | 2.2 |
| Panobinostat | 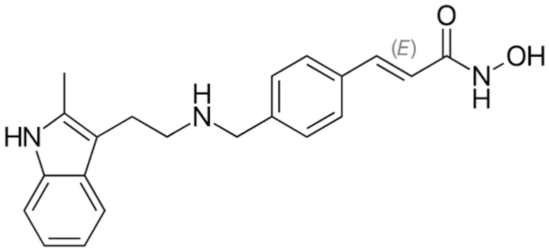 | 9.7 |

| | | |
|---|---|---|
| Fenamic Acid | 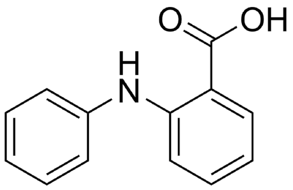 | N/a |
| Flufenamic Acid | 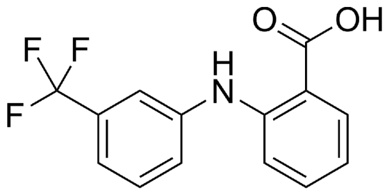 | N/a |
| Mefenamic Acid | 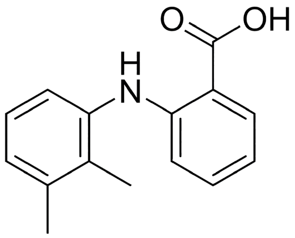 | 31.7 |
| Tolfenamic Acid | 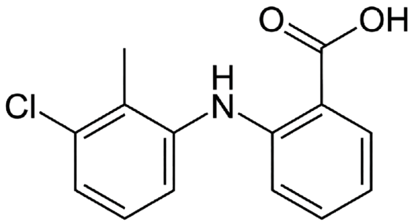 | 67.2 |
| Prulifloxacin | 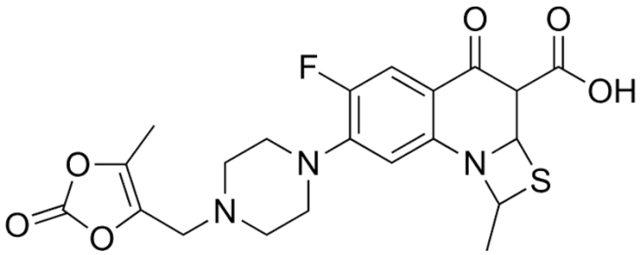 | 2.8 |

**Table S2: Boltz-2 prediction confidence.** (1) represents the overall confidence of the prediction; (2) represents the confidence in the overall structure; (3) represents the confidence in the positioning of different chains and molecules; (4) represents the confidence specifically in the model's protein-ligand interaction.[30]

| Metric | Value |
|---|---|
| 1. confidence_score | 0.903 |
| 2. ptm | 0.937 |
| 3. iptm | 0.931 |
| 4. ligand_iptm | 0.865 |

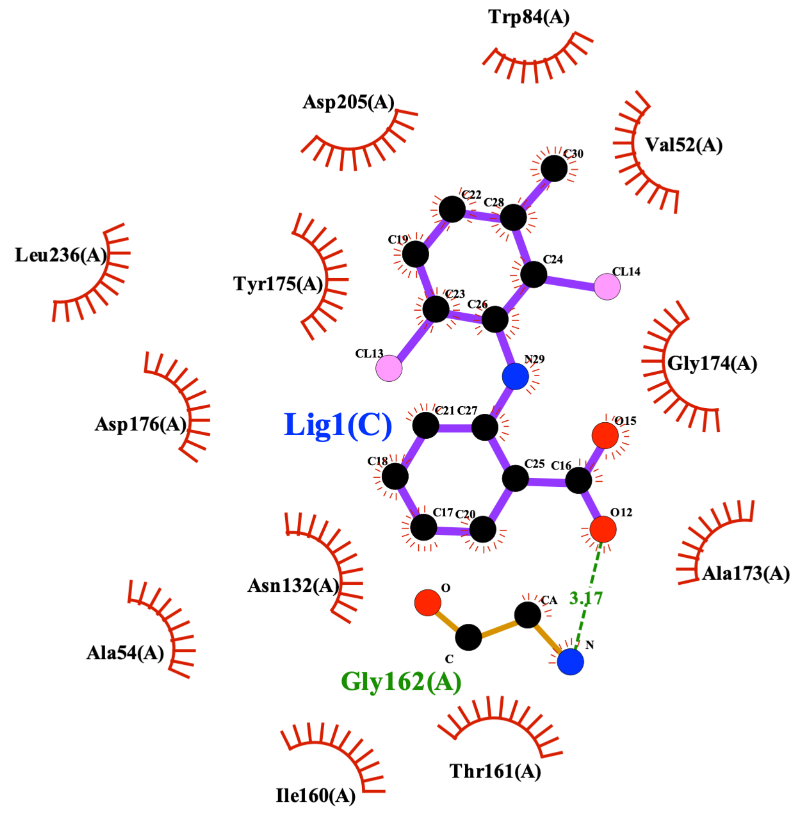


**Figure S3: Ligand-protein interaction diagram based on Boltz-2 prediction.**

```
T-COFFEE, Version_11.00 (Version_11.00)
Cedric Notredame
SCORE=97
*
 BAD AVG GOOD
*
SUGCT    :  98
FCoAT    :  97
cons     :   9

SUGCT         1 GPGSPQ---SDMNNIKPLEGVKILDLTRVLAGPFATMNLGDLGAEVIKVERPGAG-DDTRTWGPPF    62
FCoAT         1 -----MAHHHHHHA-LPLAGIKVLDLSRVLAGPWATMSLADMGAEVWKIENI-QGGDDTRAWSVP-    58

cons          1           . :   ** *:*:***:******:***.*.*:**** *:*.   * ****:*. *     66


SUGCT        63 VGTEST-YYLSV-NRNKKSIAVNIKDPKGVKIIKELAAVCDVFVENYVPGKLSAMGLGYEDIDEIA   126
FCoAT        59 NYKGASTYF-LCANRGKKSLALDLKSREGLEIIHELAKQADVVVENFRSGTVERLKIDYESLKALN   123

cons         67   . :: *:    **.***:*:::*. :*::**:***  .**.***: .*.:. : :.**.:. :    132


SUGCT       127 PHIIYCSITGYGQTGPISQRAGYDAVASAVSGLMHITG-PENGDPVRPGVAMTDLATGLYAYGAIM   191
FCoAT       124 PGIVYCSISGYGQTGPEAQRPGYDFVVQAESGLMSITGQIDG-EPTRIGVAMTDIVAGM---VATQ   185

cons        133 * *:****:******* :**.*** *..* **** ***  :. :*.* ******:.:*:    *     198


SUGCT       192 A---GLIQKYKTGKGLFIDCNLLSSQVACLSHIAAN----YLIGQKEAKRWGTAHGSIVPYQAFKT   250
FCoAT       186 SVLAALYQRKTTGLGQYIDVSLYE----CALNTLINVGSAHLNGGHVPARFGNAHPTVVPYQIFEC   247

cons        199 :   .* *: .** * :** .* .    *  :   *    :* * : . *:*.** ::**** *:    264


SUGCT       251 KDGYIVVGAGNNQQFATV-CKILDLPELIDNSKYKTNHLRVHNRKELIKILSERFEEELTSKWLYL   315
FCoAT       248 SDGAFALAVGNDRQFAILCERIIDLPELAADERFKTASGRALNRAALIPPMAER-FRTNTRQ-HWM   311

cons        265 .** :.:..**::*** :  :*:*****  :.::**   *. **  **  ::**  .  * :  ::   330


SUGCT       316 FEGS--GVPYGPINNMKNVFAEPQVLHNGLVMEME----HPTVGKISVPG-----------PAVRY   364
FCoAT       312 SACLKMGVPAGQVKTVPEAFESPNVKARQVVQKLESAHLGPISLVRPAQGLKAQENAAYKAPPMLG   377

cons        331       *** * ::.: :.* .*:*  . :* ::*     *     .. *           *.:     396


SUGCT       365 SK--------FKMSEARPPPLLGQHTTHILKEVLRYDDRAIGELLSAGVVDQHETH   412
FCoAT       378 EDSASVLGDVLGL-DGN--KL-ADLIAA--------GVIY--QYQP----------   409

cons        397 ..        : : :..   * .:  :         .     :  .             452
```

**Figure S4: Sequence alignments of human succinyl-CoA:glutarate-CoA transferase (SUGCT)[3] and formyl-coenzyme A transferase (FCoAT) from *Brucella melitensis*[18].** Multiple sequence alignment performed via T-Coffee[37].

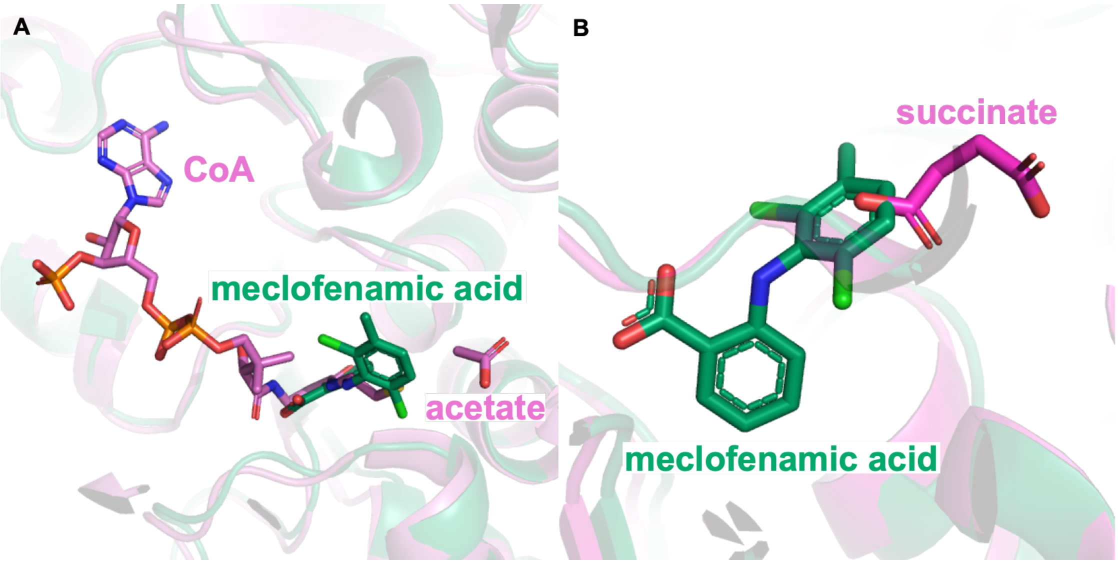


**Figure S5: Structural alignments of Boltz-2 prediction of meclofenamic acid bound to SUGCT and Formyl-coenzyme A transferase from *Brucella melitensis.*** (A) Alignment of Boltz-2 prediction (green) and PDB: 9ZAV[18] (pink). (B) Alignment of Boltz-2 prediction (green) and 10UH[19] (pink).

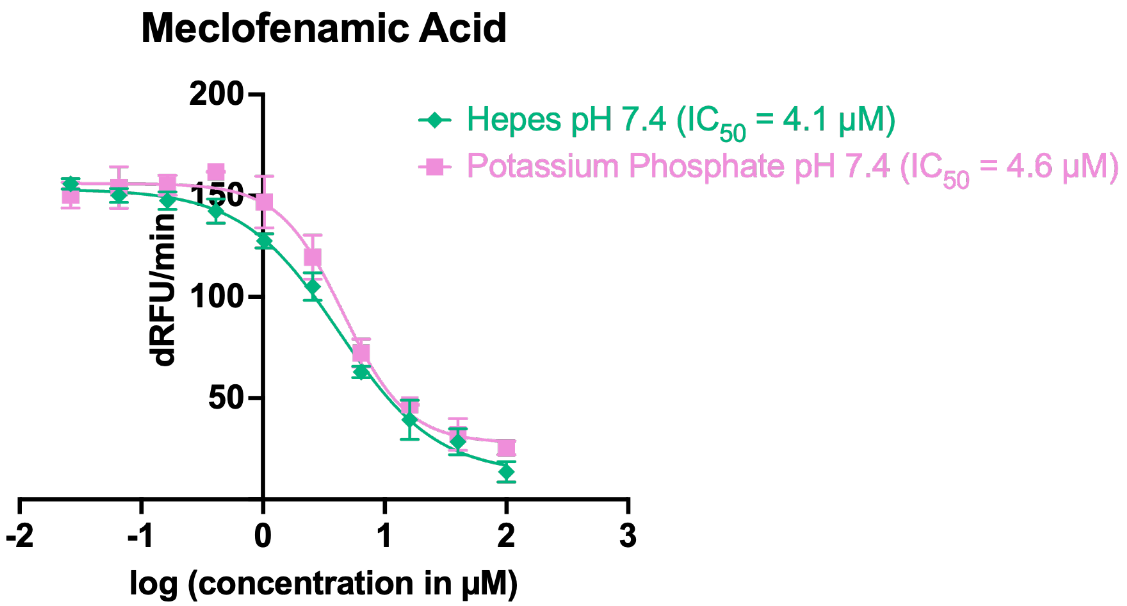


**Figure S6: Dose-response curve for meclofenamic acid in the presence of different ions.**

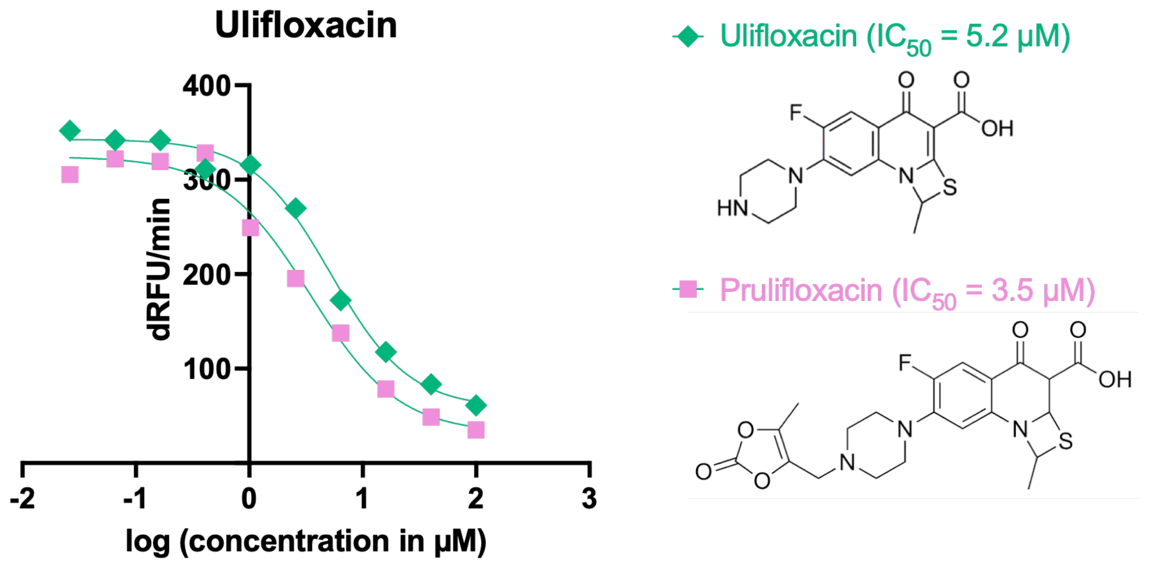


**Figure S7: Dose-response curve for ulifloxacin.**

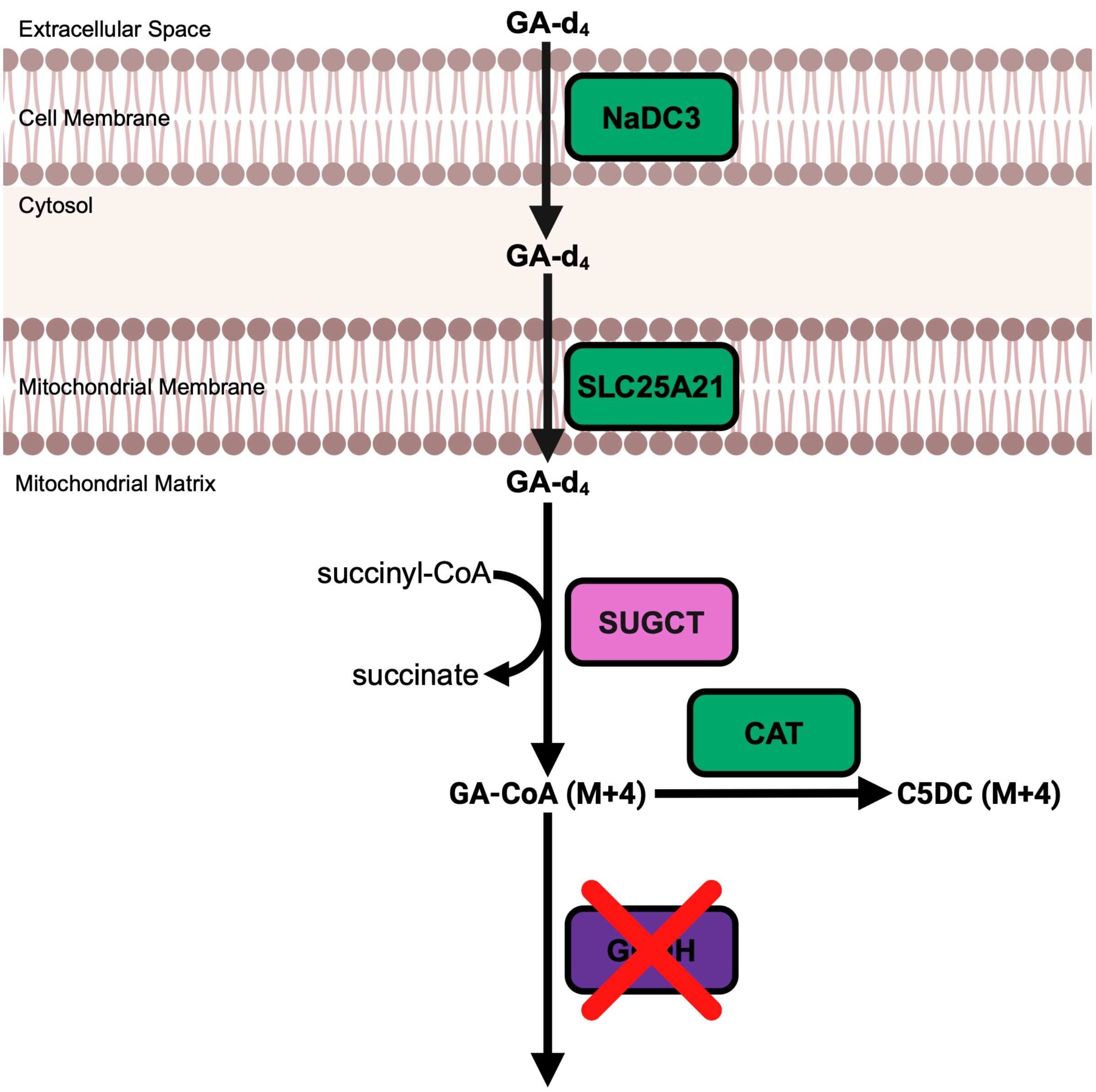


**Figure S8: Schema of cellular assay for SUGCT activity using stable isotope-labeled glutarate (GA-$d_4$).** Glutarate (GA) is a transported substrate of both the sodium-dependent dicarboxylate transporter NaDC3[24] (which is essential for cellular uptake of TCA cycle intermediates, including glutarate and succinate[25]) and the mitochondrial 2-oxodicarboxylate carrier SLC25A21[26]. A carnitine acyl transferase (CAT) mediates the conversion of glutaryl-CoA to C5DC, which can then cross membranes.[27,28]
